# Exploring the Neural Processes behind Narrative Engagement: An EEG Study

**DOI:** 10.1101/2022.11.28.518174

**Authors:** Hossein Dini, Aline Simonetti, Luis Emilio Bruni

**Affiliations:** Aalborg University, The Augmented Cognition Lab, Copenhagen, 2450, Denmark; University of Valencia, Department of Marketing and Market Research, Valencia, 46022, Spain

## Abstract

Past cognitive neuroscience studies using naturalistic stimuli have considered narratives holistically and focused on cognitive processes. In this study, we incorporated the narrative structure—the dramatic arc—as an object of investigation, to examine how engagement levels fluctuate across a narrative-aligned dramatic arc. We explored the possibility of predicting self-reported engagement ratings from neural activity and investigated the idiosyncratic effects of each phase of the dramatic arc on brain responses as well as the relationship between engagement and brain responses. We presented a movie excerpt following the six-phase narrative arc structure to female and male participants while collecting EEG signals. We then asked this group of participants to recall the excerpt, another group to segment the video based on the dramatic arc model, and a third to rate their engagement levels while watching the movie. The results showed that the self-reported engagement ratings followed the pattern of the narrative dramatic arc. Moreover, whilst EEG amplitude could not predict group-averaged engagement ratings, other features comprising dynamic inter-subject correlation, dynamic functional connectivity patterns and graph features were able to achieve this. Furthermore, neural activity in the last two phases of the dramatic arc significantly predicted engagement patterns. This study is the first to explore the cognitive processes behind the dramatic arc and its phases. By demonstrating how neural activity predicts self-reported engagement, which itself aligns with the narrative structure, this study provides insights on the interrelationships between narrative structure, neural responses, and viewer engagement.

**Significance statement:** Dramatic narratives follow a complex structure termed as the narrative arc. Here, we addressed the complexity of this structure in order to explore brain responses during narrative cognition. We examined the link between the narrative arc and its six phases with self-reported engagement, and whether brain responses elicited by a narrative can predict engagement levels. Our results showed that the group-averaged engagement ratings followed the dramatic arc model. EEG features predicted group-averaged engagement patterns and also engagement levels in the last two phases. This is the first study to characterize the narrative dramatic arc phases at the neural level. It contributes to the fields of cognitive narratology and neuroscience by extending current knowledge on how the brain responds to narratives.

## Introduction

Narratives are naturally engaging stimuli and a useful instrument for understanding affective and cognitive processes (Sonkusare et al., 2019). Regardless of their format (e.g., pictorial, written, audiovisual) or whether they are fictional or real, narratives share a general structure: beginning, middle, and end (Aristotle, 1984; Varotsis, 2018). Since Freytag’s seminal work (1895), it has been theorized that dramatic narratives follow a complex structure known as the narrative arc. A modern version of Freytag’s arc comprises six phases (the top panel of Figure 1-A): exposition, rising action, crisis, climax, falling action, and denouement (Laurel, 1991). In the exposition phase, the story’s contextual information is given. In the rising action phase, the tension created by the conflict is escalated and intensified. In the crisis phase, a dilemma related to the conflict occurs. In the climax phase, there is a turning point in which a decision is made or the dilemma is solved. From a modern perspective, the climax represents the section with the most action. In the falling action phase, the story focuses on secondary conflicts and plots. In the denouement phase, the story comes to an end. A recent study has presented quantitative evidence that narratives follow this general narrative arc (Boyd et al., 2020).

**Figure 1:**
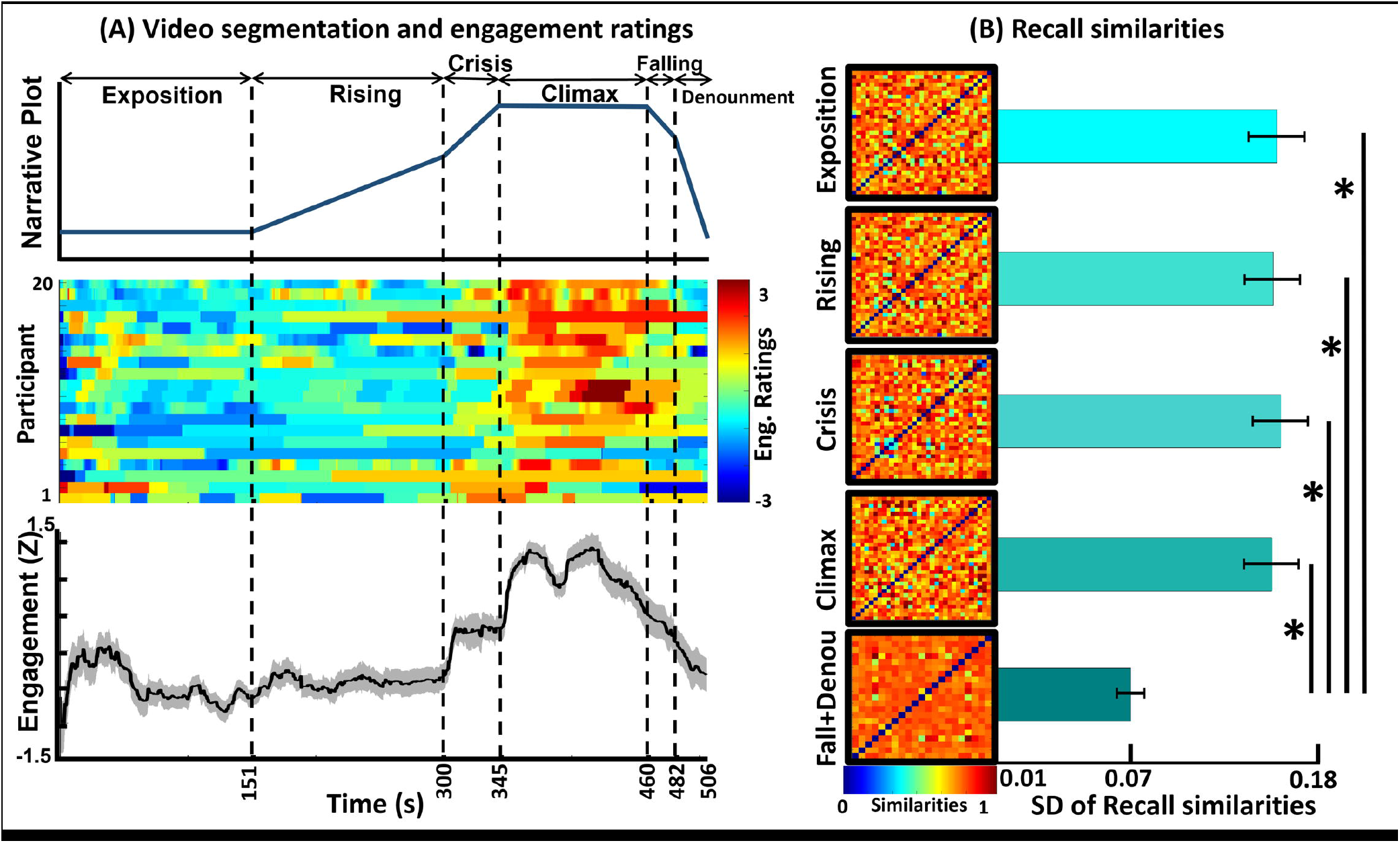
Analysis of the Segmentation, Self-reported, and Recall Parts. **A - Upper panel:** the starting point of each phase as identified by the independent participants after participating in a workshop, including the span of each phase and the phase’s names. **A - Middle panel:** the raters’ z-normalized engagement ratings. The hot colors denote higher scores, and the cold colors indicate lower scores. **A - Bottom panel:** the z-transformed group-averaged engagement ratings. The dashed vertical lines represent the start of the new phase and the end of the previous phase. **B:** the participants’ recall similarity for each separate phase. In this case, the falling action and denouement phases were combined. The matrices show the similarities between the subjects, with the hot colors indicating higher similarity scores. The horizontal bars show the standard deviations (SD) of the subjects’ recall similarities. The stars indicate that there was a significant difference (*p <* .001) between the SDs of the last phase as compared to other phases.

A narrative arc intends to create and build tension from the beginning of the story until its climax and then reduce the tension until the end of the narrative (Boyd et al., 2020). During flow state, the tension fluctuation evokes an audience’s engagement levels so that they align with the tension levels (Busselle & Bilandzic, 2009; Bilandzic et al., 2019). Engagement indeed fluctuates over the course of a narrative, and it is strong during emotional moments (Song et al., 2021b). Analyzing self-reported engagement allows for evidence of further correlation with brain responses. In fact, engaging moments in a narrative increase spectators’ shared neuronal responses (Dmochowski et al., 2012; Cohen et al., 2017; Poulsen et al., 2017; Song et al., 2021). Even though these studies used narratives as stimuli, they investigated the connection between engagement and neural responses without focusing on narrative structure. Therefore, the relationship between narrative engagement, brain responses, and narrative arc is underexplored.

Past cognitive neuroscience studies have considered narratives holistically. Hence, whether engagement levels and brain responses relate to each other in each phase of the narrative arc has not yet been investigated. The existence of a defined structure in narratives may reflect an evolutionary need for effective information sharing between individuals (Boyd et al., 2020), but an unanswered question is whether each narrative phase evokes a different brain response. The answer to this question would advance our understanding of the structure and function of narratives from a neurological perspective. In fact, the “nexus of narrative and mind” (Herman, 2009, p.30) has been the concern of cognitive narratology for the past two decades. While acknowledging the challenges of exploring the field of narrative cognition (Bruni et al., 2021), this study investigates: (a) how the engagement levels fluctuate across a dramatic arc in line with a narrative; (b) whether it is possible to predict this engagement from neural activity; and (c) in order to further examine narrative comprehension, the idiosyncratic effects of each phase of the dramatic arc on brain responses as well as the relation between engagement and brain responses are explored. We use an explorative approach to fulfill these goals. We presented an excerpt of a movie that followed the narrative arc structure to a group of participants while collecting EEG brain signals. We then asked this group of participants to recall the excerpt, another group to segment the video based on the narrative arc model, and the last group to rate their engagement levels while watching the movie. Using these datasets, we extracted several EEG features in order to explore which features better represented the link between engagement levels and brain responses in the whole narrative arc and across its phases.

## Materials and Methods

The study comprised four parts that used the same stimulus: (i) Neuro part (the EEG data); (ii) Recall part (freely recalled voice recordings); (iii) Self-reported part (self-reported engagement data); and (iv) Segmentation part (the dramatic-arc phase identification data). The data collection process for the Neuro and Recall parts was conducted in November, 2020 as part of a larger study. Participants signed an informed consent sheet prior to starting the experiment and were paid for their time. The study received approval from the ethics committee for the Technical Faculty of IT and Design at Aalborg University, and it was performed in accordance with the Danish Code of Conduct for Research and the European Code of Conduct for Research Integrity. The codes used in this study are from (Song et al., 2021a) and are addressed in the Extended data part.

### Participants, Equipment, and Experimental Design

#### NEURO AND RECALL PARTS

Thirty-two right-handed participants (13 females) with *M*_*age*_ *=* 26.84 (*SD =* 4.33, range: 20-37) wore a 32-channel EEG device (10-20 system; Brain Products). Impedance of the active electrodes was kept below the minimum threshold as stated by the manufacturer (i.e., < 25 kΩ) during the whole experiment, and the data were recorded by the Brain Products software at a sampling rate of 1,000Hz. A virtual reality headset (HTC corporation) placed on top of the EEG device was used for the stimulus presentation. We then told the participants they would see an 8 min 27 sec excerpt of an audiovisual movie, and we instructed them to pay close attention to the movie as they would be requested to recall it. The excerpt was from the movie *Pride and Prejudice and Zombies* and can be seen here: https://youtu.be/zp6EPM62wAk. We chose this subplot because it contains the six phases of the narrative arc proposed by Laurel (1991). Approximately 50 min after having watched the video, we asked the participants to recall the story from the movie excerpt aloud, and we recorded these recollections with a mobile phone.

#### SELF-REPORTED PART

Another group of 20 participants (9 females) with *M*_*age*_ *=* 24.80 (*SD =* 2.26, range: 22-30) watched the same movie excerpt on a computer. While watching it, they continuously reported their levels of their engagement by adjusting a slider scaled from 1 (“not engaging at all”) to 9 (“completely engaging”). The slider bar was constantly visible at the bottom of the screen during the experiment. This stimulus presentation and recording of responses was controlled by PsychoPy v3.0 software. Prior to the self-reported study, the definition of engagement (inspired by Song et al., 2021) was provided to the participants.

#### SEGMENTATION PART

A third group of 19 participants (8 female) with *M*_*age*_ *=* 26.10 (*SD =* 2.74, range: 23-32) attended a workshop in which they were introduced to the concept of narrative plots and the six phases of the dramatic arc structure (the top panel of Figure 1-A). Next, we asked them to watch the movie excerpt and time stamp the moments at which each of the phases started and ended in the movie.

### Data Pre-Processing

#### NEURO PART

The EEG signals were first passed through a third-order Butterworth filter that had 1 to 40 Hz cut-off frequencies to remove low and high frequency noises. After this, bad channels were detected and removed using an automated rejection procedure with a voltage threshold of ±500 _μv_. The rejected channels were then interpolated by the spherical spline method using the information from six surrounding channels in FieldTrip toolbox. The average number of rejected channels per participant was 0.93 ± 0.60. One participant was excluded due to having more than four bad channels. To remove eye-related artifacts and other remaining noises, the filtered data were fed into an independent component analysis (ICA) algorithm. Using the second-order blind identification method, the source activities (components) were estimated, and eye-related artifacts and other noise sources were detected and removed from the components list. The average number of rejected components per participant was 5.93 ± 0.97. Using the calculated ICA coefficients, the data were turned back from source space to channel space, and the de-noised data were obtained. Afterwards, the de-noised data were re-referenced to average activity of all the channels. Since a major part of the current study was based on functional connectivity analysis, an obstacle was the presence of volume conduction, which causes fake connectivity values among EEG channels (Khadem & Hossein-Zadeh, 2013). These fake connectivities exist because channels are far from source activities, so the activity of each neural source is picked up by multiple channels. To reduce the volume conduction effect, we used the current source density (CSD) method (Mitzdorf, 1985). This method obtains the spatial properties of each de-noised channel while neglecting the effect of other channels by utilizing the second spatial derivative of the EEG recorded in that channel (Dini et al., 2020). The CSD toolbox (Sijmen Duineveld, 2022) was used to apply the CSD method to the de-noised data with the medium spline flexibility of m = 4, as delineated by Dini et al. (2020) and Fitzgibbon et al. (2015). Finally, the data obtained after the implementation of CSD were z-normalized across time and down-sampled to 200 Hz to reduce the calculation load. This pre-processing procedure is demonstrated in Figure 2-A.

**Figure 2.**
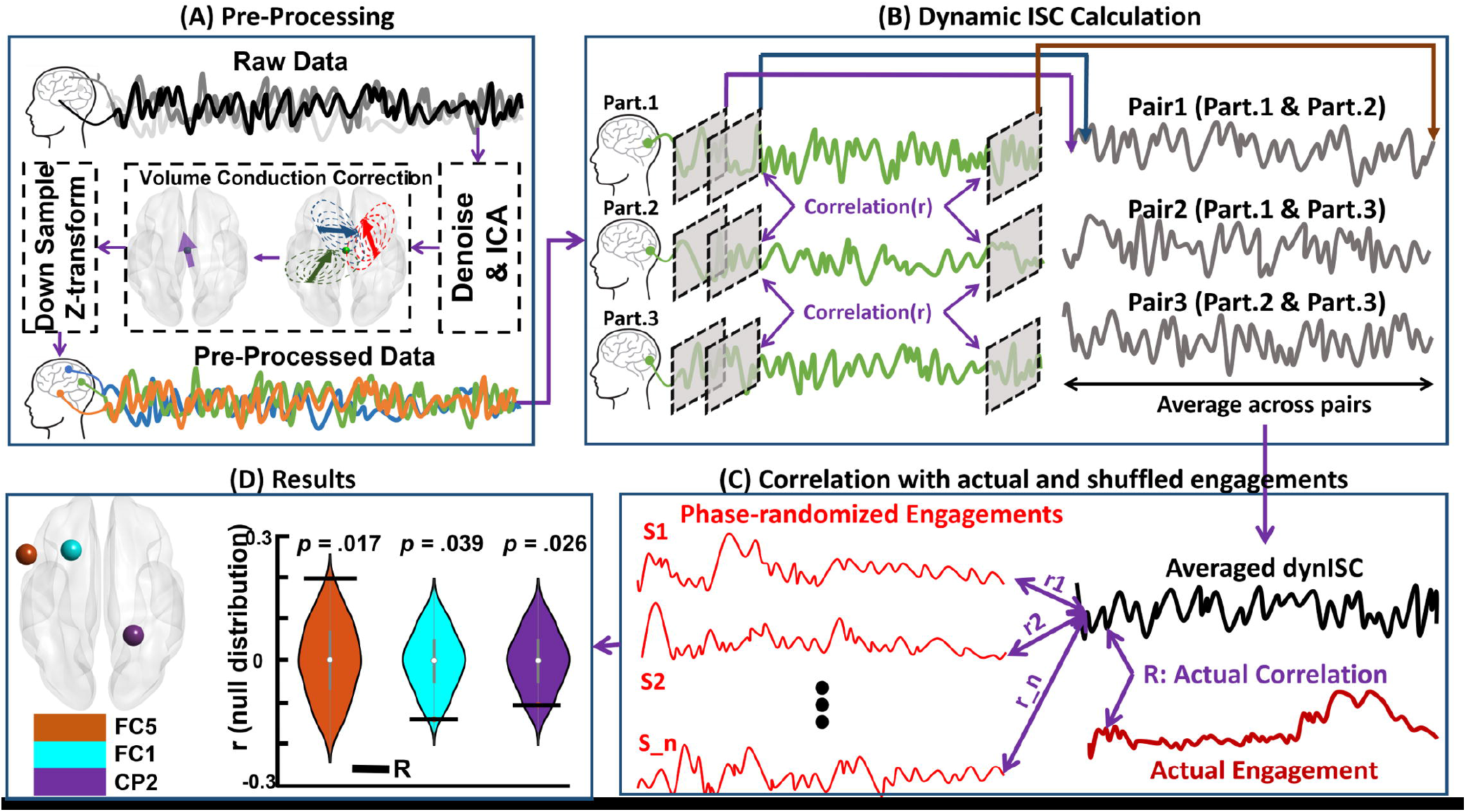
The procedure for calculating the dynamic inter-subject correlation (dISC) and the results thereof. **A:** the steps in pre-processing the EEG data. Each participant’s data went through a pre-processing stage in which the raw data were de-noised. After this, the volume conduction effect was corrected by implementing the current source density approach and then down-sampling and z-transforming. **B:** the steps for calculating the dISC for three participants (from a total of 32) and one channel (out of all the channels). The defined window (indicated by the gray) was applied to every channel for all participants. After this, we calculated the correlation between the participants’ corresponding channels for all participant pairs. Thus, this figure demonstrates the correlations for all three participant pairs as an example. The correlation between each window pair is one sample point of pairwise correlations (shown by the purple, blue, and brown arrows at the top of panel). Sliding the window throughout the entire signal, we obtained a correlation signal for the three participant pairs. The average for these correlation signals was then used in the next step. **C:** the procedure for the statistical analysis that was conducted using a permutation test. Firstly, we calculated the actual correlation between the averaged dISC and group-averaged engagement rates. Then, we phase-randomized the group-averaged correlations 10,000 times and calculated the correlation between the averaged dISC and each of these phase randomized engagement ratings. Thus, a null distribution and an actual correlation was obtained. **D:** the results from the comparison of the actual correlation to the null distribution. The orange, cyan, and, purple violin plots correspond to the FC5, FC1, and CP2 channels, respectively. The location and the obtained p-values are reported in this panel. The black horizontal lines represent the observed correlations.

### Statistical Analyses - Data Processing

In this section, we detail the methods and the implemented statistics. To facilitate easy reading, we have provided the results of the Self-reported and Segmentation parts here together with the methodology employed to obtain them.

#### SEGMENTATION PART

As previously mentioned, 19 participants attended a workshop in which they were asked to indicate the times at which each phase of the dramatic arc started after they were provided with the definitions for each phase. Using the indicated times, we explored the moments that participants consistently mentioned as the start of each phase. Implementing Silva et al.’s (2019) method, the number of participants’ indications that were different from the chance level was calculated using 3s as a window of coincidence (Baldassano et al., 2017). Shuffling the number of observations 1,000 times, a null distribution was generated. Then, the coinciding time points with a significance threshold of *p <* .050 were tested. At least eight of the participants (42.11%) should have coinciding answers that could not be explained by chance (for more on choosing the starting time points, see Silva et al., 2019). This approach resulted in the data detailed in Table 1. The dramatic arc presented by the participants is illustrated in the upper panel of Figure 1-A.

#### SELF-REPORTED PART

We then evaluated how engagement patterns changed across time as the participants experienced the story. All of the participants’ button presses were recorded and re-sampled to 200 Hz to match them with the EEG data for further correlation analysis. Continuous engagement ratings that were the same length as the video were obtained and then z-normalized across time. The participants’ engagement ratings are demonstrated in the middle panel of Figure 1-A. After this, we analyzed whether the engagement patterns were synchronized across the subjects. To do so, we calculated the pairwise correlation across engagement ratings using the ratings of all the possible participant pairs (Song et al., 2021b). The results indicate that there was a significant positive correlation between pairwise engagement ratings (mean Pearson’s *r* (18) *=* 0.40 ± 0.20). We calculated the average r by z-transforming all pairwise Pearson’s correlation r values to z space using Fisher’s method, calculating the average z values, and then transforming the averaged z value back to r value. The correlations were significantly positive in 89.47% of the pairwise correlations (false discovery rates were corrected for number of statistical tests; corrected *p <* .050). Since the participants’ engagement ratings were significantly correlated, the group-averaged engagement rating was considered representative of stimulus-related engagement (the bottom panel of Figure 1-A). A qualitative evaluation of the group-averaged engagements revealed that this engagement followed the dramatic arc pattern (e.g., engagement peaks in the climax phase and gradually decreases in the last phases).

#### NEURO PART

##### Dynamic Inter-Subject Correlation

To investigate whether neural brain activity patterns were modulated by engagement, we used dynamic inter-subject correlation (dISC). ISC is a data-driven method that correlates individuals’ neural data to others’ neural data to test whether participants perceive the same stimulus in similar ways and at the same time while neglecting subjective differences (Nastase et al., 2019). ISC is an approach that is well suited to both EEG and fMRI datasets as it captures the stimulus-driven patterns of neural activity (Simony et al., 2016; Petroni et al., 2018; Finn et al., 2020; Imhof et al., 2020; Dini et al., 2022).

Moreover, to identify the brain regions’ patterns that are modulated by stimuli, ISC is considered an alternative method that overcomes the problems of traditional methods (e.g., general linear model) as it reduces intrinsic noises (Simony et al., 2016; Song et al., 2021b). Compared to traditional methods, ISC does not require stimulus repetition and fixed experimental manipulation, which leads to a more naturalistic experimental design (Nastase et al., 2019; Song et al., 2021; Dini et al., 2022). To test the hypothesis that ISC is modulated by engagement (i.e., ISC is higher in the more engaging moments), a tapered sliding window with 15 samples (70 ms) was obtained by convolving a rectangular window with a gaussian (sigma = 3). The window size was chosen based on studies that suggest that the optimal window size for dynamic connectivity analysis is 0.05 to 0.07 multiplied by the sampling rate (Dini et al., 2021; Sendi, Zendehrouh, Sui et al., 2021; Sendi, Zendehrouh, Turner et al., 2021). Then, the designed window was slided on the pre-processed EEG signals to cover the entire duration of the signal, with the step size of 5 ms. Next, within each window, we calculated the Fisher’s z-transformed Pearson’s correlation across corresponding participant channels (e.g., correlation between the first participant’s first channel and the second participant’s first channel) and obtained the ISC for all participant pairs. Repeating the same procedure for the entire signal, we obtained the dISC for all subject pairs. The dISC calculation procedure is demonstrated in Figure 2-B. To test whether dISC was modulated by engagement, we averaged the participants’ calculated dISC according to the EEG channels.

Then, we calculated the correlation of each channel’s dISC with the group-averaged engagement ratings (see the “Self-reported part” section above) and obtained the actual correlation. To test the statistical significance of the calculated correlation, we used a permutation test. To generate the null distribution, the self-reported engagement ratings were phase-randomized, and the correlation between the calculated dISC and phase-randomized engagements were calculated for each iteration. This null distribution was generated because the phase randomization retained the characteristics of the temporal dynamic, such as frequency and amplitude, but it changed the phase (Song et al., 2021b). Repeating this procedure 1,000 times, we obtained a null distribution and tested the significance of the actual correlation assuming a one-tailed significance test with *p =* (1 + number of null r values ≥ empirical r) / (1 + number of permutations), R^2^, and the mean squared error (MSE) (find more details in Song et al., 2021). Figure 2-C demonstrates the permutation test procedure.

##### EEG Amplitude

Changes in cognitive and attentional patterns during task performance can be predicted from multivariate fMRI activity (Debettencourt et al., 2015; Haynes, 2015). Although the data used in previous studies are mainly obtained from fMRI, it was worthy to test whether it is possible to predict the engagement patterns using pre-processed EEG amplitudes without extracting any features. To do this, we applied a window with the same characteristics as above and placed the signal, and we then calculated the average EEG amplitude within each window. To predict the engagement ratings with an EEG signal, a leave-one-subject-out (LOO) cross-validation method was implemented using nonlinear support vector regression (SVR) models. The SVR models were trained using the participant’s EEG activity in each window, with one participant’s data being excluded. The SVR models were tested on the held-out participant’s EEG activity in the corresponding window to predict the group-averaged engagement ratings (see the “Self-reported part” section above). The LOO procedure is illustrated in Figure 3-D. In each cross-validation fold, the Fisher’s z-transformed Pearson’s correlation between the observed and predicted engagement ratings was calculated to serve as an indicator of predictive performance. We used this metric to evaluate the model’s performance as the main question of this study is whether the temporal dynamic patterns could be captured by the model rather than whether the model could predict the actual values of group-averaged engagement ratings. To test the statistical significance, we used a permutation test. The null distribution was generated by training and testing the same SVR models using the data from the actual brain patterns to predict phase-randomized group-averaged engagement ratings, which were repeated 1,000 times. After this, we calculated the prediction accuracy using the abovementioned correlation analysis and generated the null distribution of performance. Finally, we compared the actual model’s performance to the null distribution, assuming a one-tailed significance test with *p =* (1 + number of null r values ≥ empirical r) / (1 + number of permutations), R^2^, and MSE (see Song et al., 2021).

**Figure 3:**
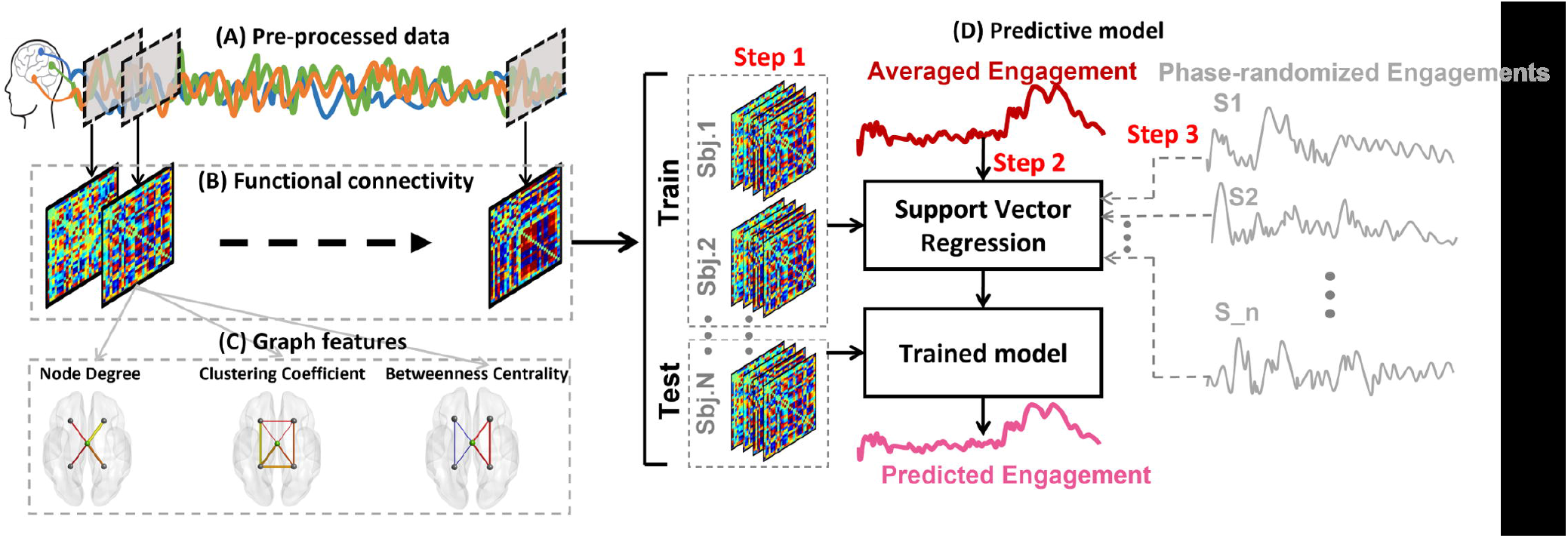
The schematic representation of the dynamic predictive model. **A:** the sample of preprocessed signals and sliding window (gray squares). **B:** the functional connectivity among all channels was obtained from the signals occurred within each window. **C:** out of each functional connectivity matrix, three graph features—node degree, clustering coefficient, and betweenness centrality—were extracted. **D:** the procedure of the predictive model. In “Step 1,” we concatenated all the functional connectivity matrices (or graph features) of all subjects and divided it to a train and test sets (the leave-one-subject-out procedure). In “Step 2,” we trained a support vector regression model using the functional connectivities of all subjects but one to predict the averaged engagements and tested the model on the leaved-out subject. The output of Step 2 is the predicted engagement rating. By calculating the correlation between actual engagement and predicted engagement, we obtained the actual correlation. In “Step 3,” we shuffled the engagement ratings as presented in Figure 2 and repeated Step 2 to predict these shuffled ratings. By calculating the correlation between the predicted engagement and each shuffled engagement scores and repeating it for all the 1,000 shuffled engagement scores, we obtained a null distribution of correlations. By testing the actual correlation on this null distribution, we obtained the significancy level of the prediction. Note that we implemented this procedure for functional connectivity features and all three graph features separately.

##### Dynamic Functional Connectivity

Changes in statistical dependence among different brain regions (i.e., functional network connectivity) could be a helpful approach to predicting changes in participants’ cognitive and attentional states while performing tasks (Rosenberg et al., 2020; Song et al., 2020; Song & Rosenberg, 2021). To calculate dynamic functional connectivity (dFC), we used the window with the same characteristics as above and placed it to cover the entire signal (Figure 3-A). Then, within each window, we calculated the Fisher’s z-transformed Pearson’s correlation across all the time periods of all channels for a single subject (e.g., the correlation between the first channel of the first subject and the second channel of the same subject). The functional connectivity matrices had a size of 32×32 to correspond with the channel numbers. This procedure is delineated in Figure 3-B. Furthermore, to test whether dFC patterns could be used to predict the engagement ratings, we used the same LOO cross-validation approach as used in the “Neuro part” investigation. The dynamic of each cell of the 32×32 matrices during the time course was considered a feature (32×31/2 = 496 features). We trained the SVR models using selected features of all participants but one, and we tested the SVR models on the selected features of the held-out participant. We employed a feature selection step prior to training the model in each cross-validation fold. The features that significantly correlated with group-averaged engagement ratings (one-sample student’s t-test with a significance threshold of *p <* .010) were selected as candidate features to train the model (Shen et al., 2017). We evaluated the predictive model’s performance using the same approach as above.

##### Graph Features

Graph-theory-based methods are helpful tools for understanding brain functional connectivity architecture (Farahani et al., 2019), and they enable better characterization of the behavior of EEG signals that simple linear methods fail to explain (Ismail & Karwowski, 2020). Therefore, it was worthy to explore how dFC structures are related to engagement prediction. To do so, we calculated three graph features using the connectivity matrices obtained from each window throughout the entire signa (Figure 3-C)l. This approach enabled us to evaluate the changes in functional connectivity structures in the time under investigation. Each electrode was considered a node, and the weighted number of correlations between the nodes were considered edges. The calculated graph features consisted of: i) a node degree (ND), which was obtained from totaling all the weights connected to a node; ii) a clustering coefficient (CC) of a node, which was determined by averaging the weights between the corresponding node and two other nodes that made a triangle with that node; iii) a betweenness centrality (BC) of a node, which was identified by calculating the number of shortest paths in the network that includes the corresponding node. The detailed definitions of the graph features selected in this study can be seen in García-Prieto et al. (2017). We extracted all the graph features using FastFC toolbox (https://github.com/juangpc/FastFC).

#### RECALL PART

We manually transcribed the full audio files recorded in the Recall part. We cleaned the transcripts by removing unrelated sentences and words (e.g., “if I remember correctly,” “what did he say,” “hmm,” etc.) for further analysis. We then segmented each participant’s transcribed recalls into six different phases based on the movie moments identified by the group of participants in the “Segmentation part” section. That is, we assigned each sentence to one of the six phases based on whether the events described in the sentence occurred within the phase’s time period. To evaluate the similarities in the participants’ recalls in each phase, we used latent semantic analysis (LSA; Landauer et al., 2013; Nguyen et al., 2019). LSA is a statistical method used to represent text similarity in semantic spaces and that has human-like performance (Landauer et al., 1998). We combined all the pre-processed recalls (text-type documents) from each phase for all participants (thirty-two) in a single big document. The LSA method was then used to identify every word in the big document and to remove repetitive and infrequent words. Next, the main components of the big document were extracted by applying a singular value decomposition method. We set the number of components to 20, which was the optimal number of components for each phase with the least perplexity that was suggested by the linear discriminant analysis (LDA) model that searched through 5, 10, 15, and 20 candidate components. As an output, the method decomposed the existing words to the bags of words (i.e., components) with similar meanings. After this, it assigned scores to each participant’s document based on their relationship with the extracted components. Finally, by calculating the pairwise distance (i.e., cosine distance) of the participant’s documents, we obtained the content similarity between all possible pairs contained in participant’s recalls.

## Results

This section presents the results obtained from the methods described in the “Neuro part” and “Recall part” in the “Materials and Methods” section.

### Recall Part

#### Participants recall the information of the falling action and denouement phases in more similar ways than in other phases

As mentioned in the Materials and Methods section, we segmented the transcribed recalls and assigned each segment to one of the phases. The length of the falling action and denouement phases was short compared to the other phases, so we merged the recall documents of those phases to ensure robust statistics. The averages and standard deviations (SD) for the number of words in each phase are as follows: exposition (*M =* 25.15, *SD =* 11.22); rising action (*M =* 17.19, *SD =* 8.34); crisis (*M =* 11.36, *SD =* 7.26); climax (*M =* 16.90, *SD =* 8.15); falling action and denouement (*M =* 13.45, *SD =* 8.94). A one-way ANOVA that had the five phases as the independent variable revealed that there was a significant difference in the number of words in the phases (*F(4, 155) =* 15.11, *p <* .001). To explore which phases differed from the others, we conducted a Dunn-Sidak post-hoc analysis. The results showed that the significant differences occurred only between the exposition phase and the other phases. This difference between the exposition phase and the other phases is expected as the exposition phase introduces the plot, main characters, and context and, therefore, contains more information than the other phases, which necessitates the explanation of more words Next, we applied LSA to the transcribed recalls of each phase and obtained the between-subjects similarity matrices as well as their SDs (Figure 1-B). Following this, we described the mean and SD of the between-subject similarities for each phase: exposition: *M =* 0.94, *SD =* 0.15; rising action: *M =* 0.94, *SD =* 0.15; crisis: *M =* 0.94, *SD =* 0.16; climax: *M =* 0.94, *SD =* 0.15; falling action and denouement: *M =* 0.97, *SD =* 0.07. As the results suggest, the falling action and denouement phase had the maximum average similarity and the lowest SD among the phases. Since we were interested in exploring how the between-subject similarities changed in each phase, we tested the statistical significance of SDs across the phases. Thus, we implemented the same one-way ANOVA as before but used the SDs obtained from each phase as the dependent variables. The results showed that there was a significant difference among the phases (*F(4, 45) =* 2.53, *p =* .015). Through a Dunn-Sidak post-hoc analysis, it was revealed that this difference was due to the comparison of the SD of the falling action and denouement phase against all the other phases (*p <* 0.05), where this phase’s SD was the lowest among all the phases. The results are illustrated as bars beside the similarity matrices in Figure 1-B. The greatest recall similarity patterns in the falling action and denouement phase implies that the participants recalled this phase in the same way (i.e., the same information was recalled).

### Neuro Part

#### Cross-subject neural synchrony follows the pattern of the narrative dramatic arc

As discussed in the “Materials and Methods” section, we calculated the dISC of the EEG signals, which served as an indicator of subjects’ neural synchrony when exposed to the narrative dramatic arc. After this, we tested our predictions of the group-averaged engagement ratings (Self-reported part) using dISC patterns. We implemented a LOO cross-validation method followed by a permutation test to compare the actual correlation with the null distribution. The results revealed that the dISC significantly predicted the group-averaged engagement ratings (two-tailed nonparametric test, *p <* .050). The dISCs of three electrodes were significantly correlated with group-averaged engagement ratings: FC5 (*r =* .20, *p =* .017); FC1 (r = -.14, *p =* .039); CP2 (*r =* -.11, *p =* .026). Out of these three channels, FC5 was positively correlated with engagement ratings, meaning that cross-subject neural synchrony increased in the more engaging moments of the narrative and decreased in the less engaging moments of the narrative. FC1 and CP2 were negatively correlated with engagement ratings, meaning that the cross-subject neural synchrony decreased in the narrative’s more engaging moments and increased in the narrative’s less engaging moments. The segmentation results and the engagement ratings indicate that the most engaging moment of the excerpt was contained within the climax phase, whereas the less engaging moments coincided with the other phases. FC5 showed the highest synchrony among the three significantly correlated channels. Two of these channels were concentrated in the frontal-central lobe of the left hemisphere (FC5 and FC1). Therefore, the results suggest that the dISC follows the narrative dramatic arc pattern in the regions concentrated in the left hemisphere frontal lobe. The results are demonstrated in Figure 2-D.

#### EEG amplitude did not provide sufficient information to predict group-averaged engagement ratings

We tested whether changes in group-averaged engagement ratings could be predicted using pre-processed EEG amplitudes. The models trained and tested based on EEG amplitudes did not result in significant predictions of group-averaged engagement ratings (*p =* .521, *r =* - .015, *MSE =* 1.064, *R*^2^ *=* -.064) in any of the EEG channels. The reported values were the result of a correlation analysis between predicted engagements based on EEG amplitudes and actual group-averaged engagement ratings that were averaged across the channels. Furthermore, we added a feature selection section in every cross-validation iteration to increase the neural features’ specificity. Therefore, only the EEG time courses (channels) that were consistently correlated with the group-averaged engagement ratings (one-sample student’s t-test, *p <* -.010) were fed into the SVR models instead of the data from all channels (Shen et al., 2017). Training the models using the selected features still did not result in robust predictions, and the EEG amplitudes were not significantly correlated with engagement ratings (*p =* .438, *r =* -.018, *MSE =* 1.021, *R*^2^ *=* -.061). Therefore, our results show that EEG amplitudes failed to predict the group-averaged engagement ratings. The results obtained from testing the actual prediction values on the generated null distribution is illustrated in Figure 4-A in the form of a gray violin plot.

**Figure 4.**
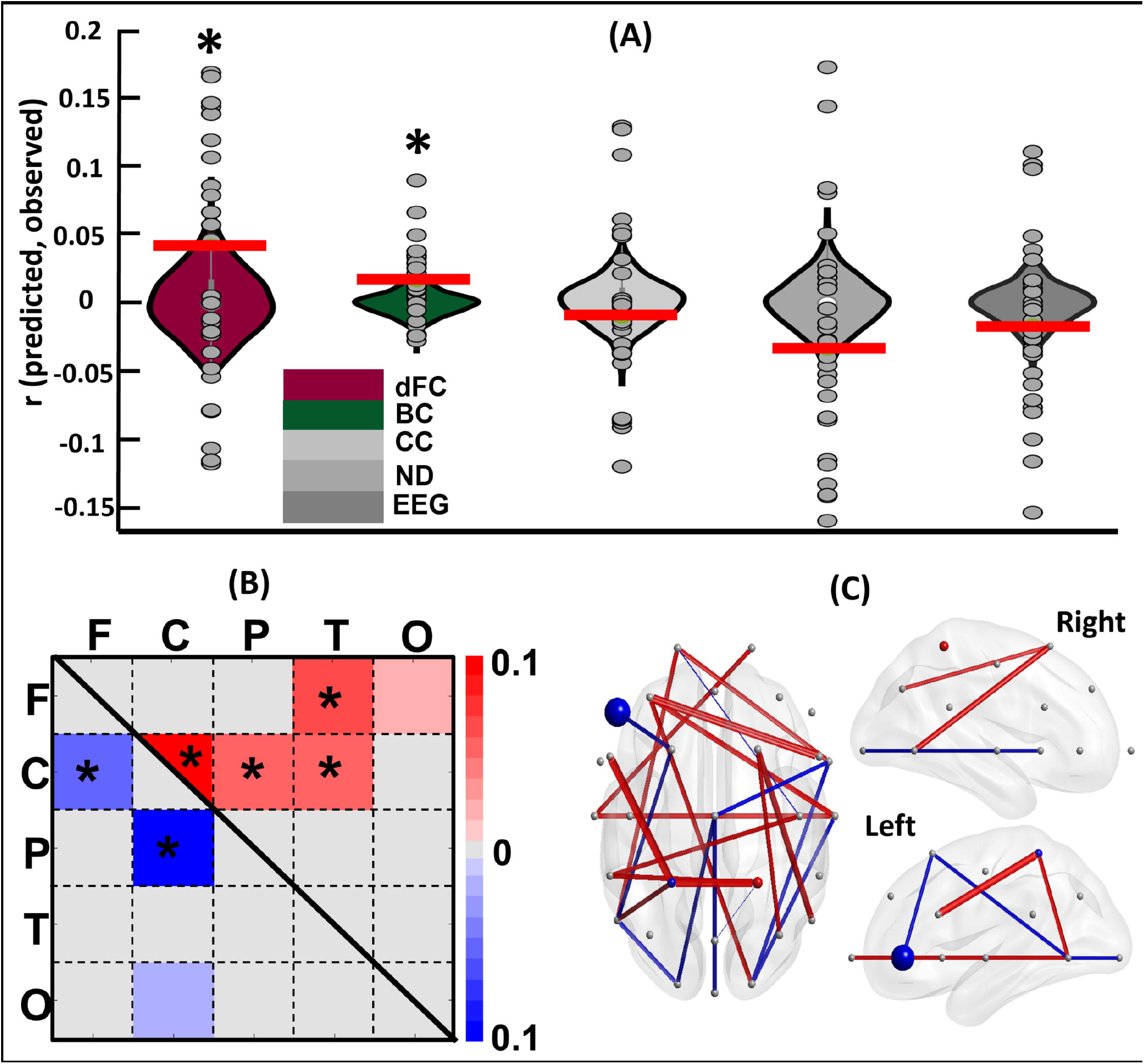
The engagement level predictions for the whole narrative. **A:** the testing of the actual correlation against the null distribution. Each violin plot refers to the prediction results for each EEG feature. The colored violin plots refer to statistically significant prediction features, and the gray plots refer to the features that were not statistically significant. The dark red violin plot refers to the dFC feature, and dark green plot refers to the BC feature. The light to dark gray spectrum refers to the CC, ND, and EEG features, respectively. The gray circles in the violin plots show the actual correlation obtained in each cross-validation fold with reference to each participant. The red horizontal lines show the observed correlation. The stars indicate that the corresponding feature had been a predictor that was chosen due to it being significantly higher than a chance occurrence. **B:** the predictor FCs that are either positively (red) or negatively (blue) correlated with engagement ratings across brain regions (F = frontal, C = central, P = parietal, T = temporal, O = occipital). Each cell represents the average number of times that each region played a predictor role. The light to dark colors (red and blue) refer to the strength of their score, which is the number of times they were considered a predictor divided by the length of all possible connections with other channels. **C:** the predictor FC features and channels on the scalp from three perspectives. The predictor FCs that were averaged in panel B are shown individually here in that the positively correlated items are indicated by red and the negatively correlated items are indicated by blue. The predictor channels obtained from the BC features are represented as colored dots, with bigger dots denoting higher proportions.

#### dFC patterns could predict the group-averaged engagement ratings in which the central region plays a vital role

The models trained and tested using extracted functional connectivity (FC) features successfully predicted the group-averaged engagement ratings (*p =* .034, *r =* .043, *MSE =* 1.115, *R*^2^ = -.115), as shown in Figure 4-A in the form of a dark red violin plot. The null distributions are positively skewed in this figure, which might be because they were generated by training the model to predict the phase-randomized engagements and because the model was tested to predict the same data using the held-out participant’s dFC. Nevertheless, the prediction accuracy was significantly higher than the generated null distribution. To investigate which brain regions significantly contributed to the predictions of group-averaged engagement, we visualized the FC features that were consistently selected as predictors (see the “Data Processing,” “Neuro Part,” and “Dynamic Functional Connectivity” sub-sections) in every cross-validation fold seen in Figure 4-B and 4-C. A total of 264 FC features that were positively correlated with engagement and a total of 136 FC features that were negatively correlated with engagement were both consistently selected. We called these features “predictive FCs.” Each participant had a matrix consisting of these positively and negatively correlated features. Figure 4-C shows the selected FC features averaged across participants, with red denoting positively correlated features and blue signaling negatively correlated features. To have a better understanding of these predictive FCs, we first defined the five canonical brain regions: frontal (F), central (C), parietal (P), temporal (T), and occipital (O). After this, we totaled the number of predictive FC features included in each region, resulting in a 5×5 matrix for each participant. Next, we calculated the proportion of predictive FCs relative to the total number of possible connections among regions to create a proportion matrix for each participant. This was done by dividing the 5×5 matrices by the lengths of all possible connections among regions (i.e., networks). Finally, by averaging the proportion matrices across the participants, we obtained the proportion score of all possible connections (see Figure 4-B).

To evaluate whether the defined networks—containing predictive FC information—were present more frequently than would be explained by chance, we implemented a one tailed non-parametric test on the number of predictive FCs within and between regions (the significant regions are indicated by stars in Figure 4-B; FDR-corrected *p <* .001). All the p-values were then corrected according to the number of test repetitions using the Benjamini-Hochberg method. Our results suggest that the within-central, between-central, and all other regions except occipital were constantly and significantly selected as predictive FC features. Although there was also a frontal-temporal connection that was able to predict the engagement pattern, our results suggest that the central region plays a vital role in predicting engagement patterns.

#### Graph features of central and frontal regions could significantly predict the group-averaged engagement ratings

We examined whether the graph features extracted from the connectivity matrices described in the previous section could predict engagement ratings. The SVR models were trained and tested using three graph features: ND, CC, and BC. The procedure for the LOO cross-validation and the subsequent permutation test was the same as described previously. The results show that BC significantly predicted the group-averaged engagement ratings (*p =* .027, *r =* -.016, *MSE =* 1.061, *R*^2^ *=* -.061). As shown in Figure 4-A in dark green, the actual correlation is significantly higher than the generated null distribution. However, ND and CC failed to significantly predict the engagement ratings (ND: *p =* .351, *r =* .007, *MSE =* 1.110, *R*^2^ *=* -.011; and CC: *p =* .756, *r =* -.009, *MSE =* 1.062, *R*^2^ *=* -.062). As shown in Figure 4-A in gray, these aspects were not significantly distanced from the null distributions. For BC, we evaluated which channels were consistently selected in each cross-validation fold. The results showed a total of three channels, with the CP2 channel being positively correlated and the CP1 and F7 channels being negatively correlated with the group-averaged engagements. In Figure 4-C, the predictive channels with positive correlations are labelled in red, and those with negative correlations are labelled in blue. Next, we evaluated whether the repetition of these channels is higher than would be expected by chance by calculating the channels’ proportion of predictive BC relative to the total number of possible channels (as above). The results show that all three channels’ BCs were significantly selected in cross-validation folds (CP2: proportion score = .066, corrected *p <* .001; F7: proportion score = .433, corrected *p <* .001; and CP1: proportion score = .033, corrected *p <* .001). Therefore, the graph theoretical feature from central and frontal regions could significantly predict the group-averaged engagement ratings.

#### The neural activity of the falling action and denouement phase could successfully predict engagement patterns in which the central and frontal regions play a vital role

We made advancements in evaluating neural activity that was stimulated by each narrative phase as opposed to just making predictions for the whole excerpt. For this, we analyzed the possibility of predicting the engagement levels in each phase separately using the data obtained from the brain. We segmented both the group-averaged engagement ratings and neural activity based on the phases delineated in the “Segmentation part” section. We extracted dFC and BC features, in the same way that we extracted to predict the whole narrative, this time separately for six phases. After this, we implemented the same LOO cross-validation approach to evaluate whether the feature could predict the engagement ratings. All the methods implemented herein were identical to those explained previously, but the models were fed the neural activity and engagement ratings of each phase, and they were trained and tested separately in this case. The results show that the models trained and tested using the dFC features of each phase failed to significantly predict the exposition (*p =* .798, *r =* -.009, *MSE =* 1.149, *R*^2^ *=* -.149), rising action (*p =* .516, *r =* .023, *MSE =* 1.569, *R*^2^ = -.569), crisis (*p =* -.687, *r =* -.018, *MSE =* 1.577, *R*^2^ = -.577), and climax (*p =* .197, *r =* -.018, *MSE* = 1.577, *R*^2^ = -.577) phases. However, the models significantly predicted the falling action (*p =* .036, *r =* -.297, *MSE =* 1.708, *R*^2^ *=* -.708) and denouement (*p <* .001, *r =* -.355, *MSE =* 1.946, *R*^2^ = -.946) phases. The results (dFC features) obtained from comparing each phase’s actual correlation with the generated null distribution are demonstrated in Figure 5-A, which shows that the actual correlation was significantly lower than the null distribution in the falling action and denouement phases. Furthermore, the results from training and testing the models that were fed the BC features of each phase revealed that, for the dFC feature, the models failed to significantly predict the exposition (*p =* .947, *r =* -.023, *MSE =* 1.039, *R*^2^ = - .039), rising action (*p =* .399, *r = -*.003, *MSE =* 1.342, *R*^2^ *=* -.342), crisis (*p =* .714, *r =* -.014, *MSE =* 3.374, *R*^2^ *=* -.374), and climax (*p =* .457, *r =* .003, *MSE =* 1.468, *R*^2^ = -.468) phases. However, the models predicted the falling action (*p =* .072, *r =* - .046, *MSE =* 1.401, *R*^2^ *=* - .401) and denouement (*p =* .065, *r =* .072, *MSE =* 1.303, *R*^2^ *=* -.303) phases only marginally significant. The results (BC features) from comparing each phase’s actual correlation with the generated null distribution are demonstrated in Figure 5-B, which shows that the actual correlation was significantly higher than the null distribution in the falling action and denouement phases.

**Figure 5:**
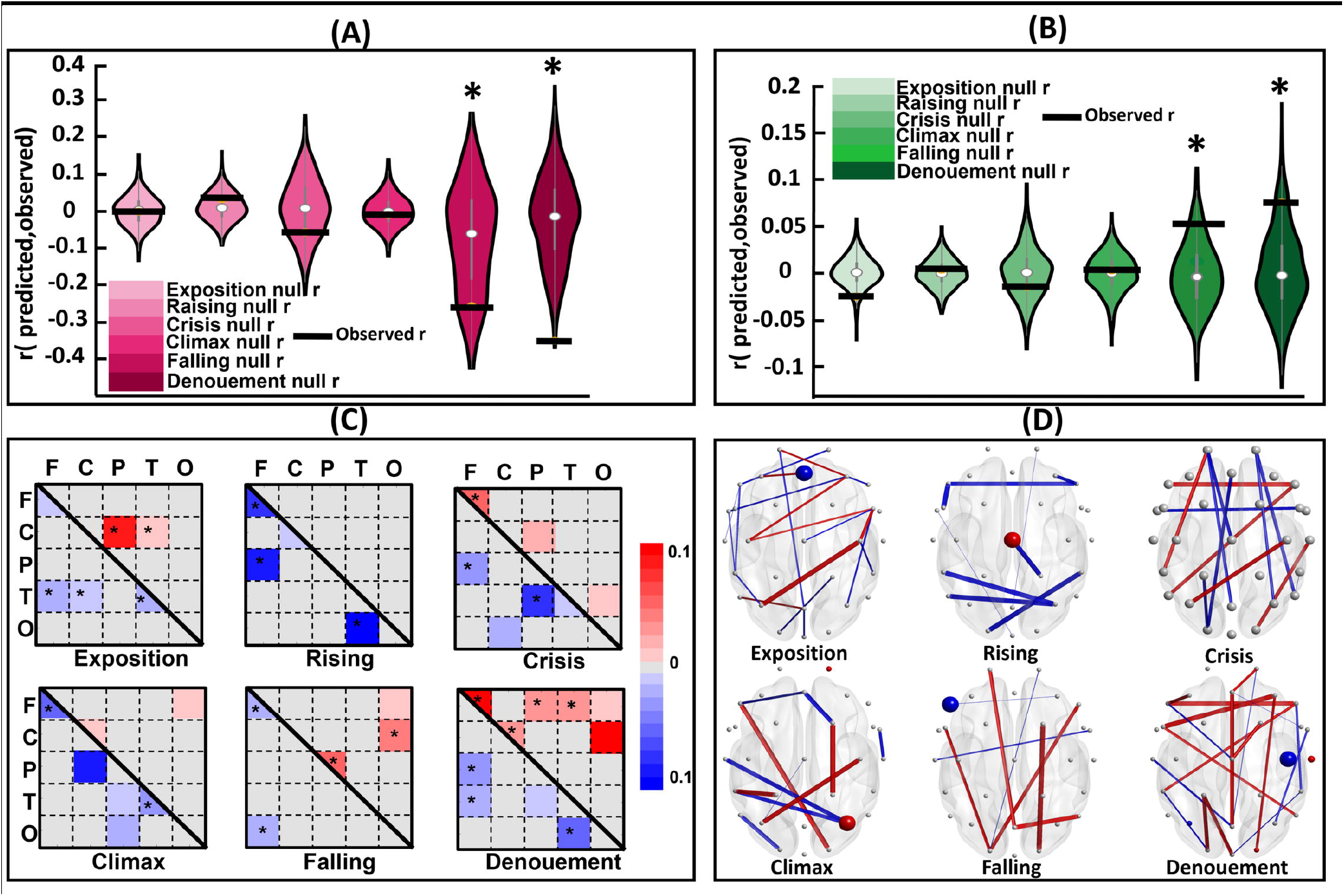
see Figure 4. The predictions of the engagement ratings of each phase using the corresponding features. The black horizontal lines show the observed correlation, and the violins show the null distribution. **A:** the phase prediction results for all phases using dFC features. It shows that the prediction in the last two phases (falling action and denouement) was significantly higher than could be explained by chance. **B:** the prediction results using BC features and the significant prediction that occurred in the last two phases. **C:** the predictor FCs across regions and their proportion scores for each phase. **D:** the predictive FCs and the channels that the BCs played as predictor roles for each phase.

The FCs that were consistently selected as predictors in each cross-validation fold are shown in Figure 4-D, with a separate label for each phase. The total number of positively and negatively correlated predictor FCs are as follows: exposition: 96 positively and 7 negatively correlated; rising action: 0 positively and 170 negatively correlated; crisis: 52 positively and 108 negatively correlated; climax: 14 positively and 130 negatively correlated; falling action: 40 positively and 32 negatively correlated; denouement: 184 positively and 96 negatively correlated. As in the previous section, we examined FCs that acted as predictors in each brain region defined above. Figure 5-C shows these FCs, with a separate label for each phase. After this, we tested whether these regions were selected in ways that were significantly higher than would be expected for chance selections by using the statistical analysis explained above. The significant predictor FCs are indicated by the stars in the corresponding matrix of each phase in Figure 5-C. The results demonstrate that, in terms of the phases in which the SVR model could successfully predict the engagement ratings (i.e., falling action and denouement), the frontal lobe and the connection of the frontal region with other regions played a vital role in predicting the engagements: In the falling action phase, the connection was between frontal and occipital, and, in the denouement phase, the connection was between frontal and parietal and temporal regions. Moreover, the central-occipital connection in the falling action phase and the central connection in the denouement phase were also significant predictors. Although the neural activity of the other phases failed to predict the engagement ratings, it is worth mentioning that, in all of them except for the exposition phase, the frontal region was identified as a significant predictor.

We then implemented the approach explained in the “Graph” section to investigate which BC of which channels were consistently selected as predictors by calculating the proportion scores. In the last two phases, in which the model successfully predicted the engagement ratings using the BC feature, the selected channels, their calculated proportion score, and the p-values were as follows: falling action: F7, negatively correlated, proportion score = .033, *p =* .029; denouement: C4, negatively correlated, proportion score = 1, *p <* .018; P3, negatively correlated, proportion score = .066, *p =* .018; O2, positively correlated, proportion score = .030, *p =* .031; T8, positively correlated, proportion score = .200, *p <* .001. Therefore, in the denouement phase, the highest-scoring channel (C4) was in the central region. Moreover, considering all of the phases, most of the predictor channels were related to the central and frontal regions.

In summary, our results show that the models trained on dFC and BC features were able to significantly predict the engagement ratings of the falling action and denouement phases. Moreover, the frontal and central regions as well as their connection to other regions played an important role in predicting the engagement ratings in the last two phases, which was determined by the evaluation of both dFC and BC features.

## Discussion

Higher levels of narrativity are ensured in stories that have a dramatic arc (Ryan, 2007). We addressed the complexity of the dramatic arc in order to explore brain responses during narrative cognition. Specifically, we examined the link between the phases of a narrative arc and self-reported engagement and whether brain responses elicited by a narrative can predict engagement levels. Our results show that the group-averaged engagements followed the dramatic arc model and could be predicted by dISC, dFC, and BC features. We also predicted engagement in the last two phases by investigating the neural underpinnings of each phase. The fluctuations in self-reported engagement ratings occurred synchronously for all the participants, reflecting that individuals share states of engagement. Moreover, individuals could identify a narrative arc structure in our stimulus (the upper panel of Figure 1-A). Notably, we found that perceived engagement at the group level followed the same shape as the narrative arc structure (the bottom panel of Figure 1-A). The individuals’ synchronous engagement and the shape of its temporal development (i.e., similar to the narrative arc) could be attributed to transportation effects (see Green & Brock, 2000). This finding is also in line with Song et al.’s (2021) study in which it was shown that emotional moments of a narrative evoke stronger engagement levels. Moreover, previous studies have reported that an individual’s engagement state is evoked by and aligned with tension levels (Busselle & Bilandzic, 2009; Bilandzic et al., 2019). Similarly, our results show that group-averaged engagement increased in the rising action phase, became more pronounced in the crisis phase, and peaked in the climax phase in which there is maximum tension, after which it declines. Therefore, the dramatic arc regulates an individual’s engagement state by fluctuating tension levels.

We then explored whether individuals’ brain activity was modulated by their engagement levels throughout the dramatic arc model. We previously found that different levels of narrativity led to differences in ISC and different levels of perceived engagement, thereby predicting narrativity level (Dini et al., 2022). Cohen et al. (2017) evaluated time perception and its relation to engagement during narrative videos, from which they posited that engagement could be considered subject’s levels of similarity among EEG channels (i.e., ISC). Moreover, the subjective perception of time and neural processing becomes synchronous when watching narratives (Cohen et al., 2017). Dmochowski et al. (2012) stated that extracting correlated components using ISC is related to “emotionally-laden attention.” Thus, ISC might be a marker of engagement (Dmochowski et al., 2012). Correspondingly, our results indicate that the dISC of individuals’ frontal (FC5 and FC1) and central (CP2) electrodes were more synchronized in a narrative’s more engaging moments. Implementing fully cross-validated models revealed that the dISC of the frontocentral region became more active and involved in predicting moment-by-moment engagement levels (Figure 5-D), whereas EEG amplitudes could not make this prediction (Figure 4-A). Therefore, the dISC patterns followed the dramatic arc structure. The patterns were organized in a way that subjects’ neural activity was synchronized in the engaging phases of the dramatic arc (i.e., crisis and climax) and relatively less synchronized in less engaging moments (i.e., exposition, rising action, falling action, and denouement). While previous studies also found a link between engagement and ISC levels (which supports the use of dISC as a marker of engagement), we extended this and argued that the subjects’ neural activity was modulated by the presence of a narrative structure that unfolded as a dramatic arc. This finding is a promising step forward in the understanding of narrative cognition.

Previous studies have also characterized engagement as well as emotional and focused attentional states, which is closely related to engagement (Busselle & Bilandzic, 2009; Dmochowski et al., 2012) using FNC. In a dFC study, Song et al. (2021) evaluated narrative engagement, sustained attention, and event memory during narrative exposure. They reported that models based on dynamic brain connections could predict engagement states in two independent datasets. Moreover, the default mode network activity, which fluctuated in line with engagement states, was also able to predict sustained attention and recall of the narrative events. In the current study, we explored if there was a link between engagement states and FNC and FNC-related features (i.e., BC graph). By using similar cross-validation models, both dFC features and the BC graph feature were significantly correlated with engagement states (Figure 4-A). The FC networks’ patterns that acted as predictors revealed that the central and frontal lobe played a vital role in predicting engagement states (Figure 4-B). Moreover, the predictor graph channels were concentrated on central (CP2, CP1) and frontal (F7) channels. In line with the findings on dISC predictors, the dFC confirmed that the neural connections across central and frontal regions were more synchronous with engagement patterns (i.e., more neural activity in more engaging moments of the narrative and vice versa). In addition, the BC graph feature, in line with the dISC (specifically in the CP2 channel), revealed that the central and frontal channels played a crucial role in predicting engagement patterns. The BC graph features have been considered putative hubs in a network (Bullmore & Sporns, 2009). In summary, the analysis of the dFC and BC features showed that predictive channels in frontal and central regions followed the dramatic arc pattern (confirmed by the dISC) and that the information flow passed (or possibly accumulated) through the regions of those channels to connect to other brain regions. Hence, these features were important for characterizing predictive brain regions for cognitive, attentional, and engagement states, in addition to ISC.

This is the first study that has characterized the narrative dramatic arc phases at the neural level. Past studies have investigated narrative phases (Bullmore & Sporns, 2009), but have not explored the neural processes that underlie the information processing of each phase. Based on machine-based semantic analysis, Boyd et al. (2020) reviewed approximately 40,000 traditional narratives and provided evidence that they consist of a start, which is followed by plot progression, and they then end with a decrease in cognitive tension. They claim that their study was the first that provided empirical support for dramatic arc narrative structures, and they suggested that future studies should focus on exploring psychological aspects induced by this structure. In this study, we used EEG to explore the neuropsychological underpinnings of engagement states for each phase. The results show that both dFC and BC could predict engagement but only for the falling action and denouement phase (Figure 5-C, D). In this case, as with the prediction of the whole narrative, the frontal and central electrodes were the most influential. The last two phases were the least complex in the excerpt used for this study: They only featured one scene with only two characters who had already been introduced, and they did not contain any background action. The higher levels (and low SDs) of the participants’ recall similarity for these phases supported the idea that these phases conveyed more straightforward and less information (Figure 1-B). Previous literature on situation models (e.g., Zwaan, 1999; Huff et al., 2014; Lin et al., 2019) state that past individual experiences and the information provided by the narrative shape the construction of situation models used for narrative interpretation. Moreover, multiversional thinking proposes the idea that multiple versions of the same narrative can coexist in the audience’s mind as the narrative progresses (Hiskes et al., 2022). We argue that, in our excerpt, the last two phases created much less opportunity for multiversional thinking and complex situation models, which might have been reflected in the pattern of brain activation. This type of brain activation was able to predict the engagement level, but the activation when facing higher levels of complexity in the narrative (i.e., increased amount of information) could not. This allows for the exploration of which EEG features could predict engagement in the latter case.

This study contributes to the fields of cognitive narratology and neuroscience by extending our (still limited) knowledge on how brains respond to narratives and how these responses are linked to perceived engagement. In addition, we extended the knowledge on using narratives not only as a means but also as objects of research by analyzing brain responses, perceived engagement, and dramatic arc structure relationships. The movie excerpt selected in this study acted as a fraction of a dramatic arc within the complete narrative (the whole movie). Although the excerpt was perceived as possessing a dramatic arc structure, it did not allow us to make links between the phases and more complex narrative features, such as expectancy and closure. Thus, future studies can advance the study of brain responses to narrative features by presenting a complete narrative to participants. Moreover, given the unique nature of each narrative, there is a need for more studies that use different narratives before our findings can be generalized (Vaccaro et al., 2021). Furthermore, an ambitious step would be investigating how brain responses differ between linear and non-linear narratives, which opens the door for the exploration of cognitive processes in interactive and immersive narratives (Bruni et al., 2022).

## Acknowledgments

This work was supported by Rhumbo (European Union’s Horizon 2020 research and innovation program under the Marie Skłodowska-Curie Grant Agreement No 813234). We are grateful to Tirdad Seifi Ala for helping with technical aspects of the data analysis and for revising an early version of the manuscript, to Thomas Anthony Pedersen and Nele Kadastik for inputs in later discussions, and to Mohammad Jafarian for helping to create the figures.

## Extended Data

The codes for FC feature extraction and the SVR model are provided by (Song et al., 2021a).The same procedure was used in this study. The sample data and the code are available at Github : https://github.com/hyssong/NarrativeEngagement.

## TABLE

**Table 1.** The starting time points indicated by the participants. The choice of the starting time point of each phase of the dramatic arc was obtained from the data of 19 participants that indicated these moments. From the starting time points, the duration of each phase was calculated. The number of participants that indicated the same time points statistically and the SD of all responses are indicated in the last two columns, respectively.

## References

Baldassano, C., Chen, J., Zadbood, A., Pillow, J. W., Hasson, U., & Norman, K. A. (2017). Discovering Event Structure in Continuous Narrative Perception and Memory. Neuron, 95(3), 709-721.e5. https://doi.org/10.1016/j.neuron.2017.06.041

Bilandzic, H., Sukalla, F., Schnell, C., Hastall, M. R., & Busselle, R. W. (2019). The narrative engageability scale: A multidimensional trait measure for the propensity to become engaged in a story. International Journal of Communication, 13, 801–832. https://ijoc.org/index.php/ijoc/article/view/8624

Boyd, R. L., Blackburn, K. G., & Pennebaker, J. W. (2020). The narrative arc: Revealing core narrative structures through text analysis. Science Advances, 6(32), 1–10. https://doi.org/10.1126/sciadv.aba2196

Bruni, L. E., Dini, H., & Simonetti, A. (2021). Narrative Cognition in Mixed Reality Systems: Towards an Empirical Framework. In J. Y. C. Chen & G. Fragomeni (Eds.), Virtual, Augmented and Mixed Reality (pp. 3–17). Springer International Publishing. https://doi.org/10.1007/978-3-030-77599-5_1

Bruni, L. E., Kadastik, N., Pedersen, T. A., & Dini, H. (2022). Digital Narratives in Extended Realities. In Roadmapping Extended Reality: Fundamentals and Applications. Wiley.

Bullmore, E., & Sporns, O. (2009). Complex brain networks: graph theoretical analysis of structural and functional systems. Nature Reviews Neuroscience, 10(3), 186–198.

Busselle, R., & Bilandzic, H. (2009). Measuring Narrative Engagement. Media Psychology, 12(4), 321–347. https://doi.org/10.1080/15213260903287259

Cohen, S. S., Henin, S., & Parra, L. C. (2017). Engaging narratives evoke similar neural activity and lead to similar time perception. Scientific Reports, 7(1), 4578. https://doi.org/10.1038/s41598-017-04402-4

Debettencourt, M. T., Cohen, J. D., Lee, R. F., Norman, K. A., & Turk-Browne, N. B. (2015). Closed-loop training of attention with real-time brain imaging. Nature Neuroscience, 18(3), 470–475.

Dini, H., Ghassemi, F., & Sendi, M. S. E. (2020). Investigation of Brain Functional Networks in Children Suffering from Attention Deficit Hyperactivity Disorder. Brain Topography, 33(6), 733–750.

Dini, H., Sendi, M. S. E., Sui, J., Fu, Z., Espinoza, R., Narr, K. L., Qi, S., Abbott, C. C., van Rooij, S. J. H., & Riva-Posse, P. (2021). Dynamic functional connectivity predicts treatment response to electroconvulsive therapy in major depressive disorder. Frontiers in Human Neuroscience.

Dini, H., Simonetti, A., Binge, E., & Bruni, L. E. (2022). Higher levels of narrativity lead to similar patterns of EEG activity across individuals. BioRxiv.

Dmochowski, J. P., Sajda, P., Dias, J., & Parra, L. C. (2012). Correlated Components of Ongoing EEG Point to Emotionally Laden Attention – A Possible Marker of Engagement? Frontiers in Human Neuroscience, 6(MAY 2012), 1–9. https://doi.org/10.3389/fnhum.2012.00112

Farahani, F. V, Karwowski, W., & Lighthall, N. R. (2019). Application of graph theory for identifying connectivity patterns in human brain networks: a systematic review. Frontiers in Neuroscience, 13, 585.

Finn, E. S., Glerean, E., Khojandi, A. Y., Nielson, D., Molfese, P. J., Handwerker, D. A., & Bandettini, P. A. (2020). Idiosynchrony: From shared responses to individual differences during naturalistic neuroimaging. NeuroImage, 215, 116828.

Fitzgibbon, S. P., DeLosAngeles, D., Lewis, T. W., Powers, D. M. W., Whitham, E. M., Willoughby, J. O., & Pope, K. J. (2015). Surface Laplacian of scalp electrical signals and independent component analysis resolve EMG contamination of electroencephalogram. International Journal of Psychophysiology, 97(3), 277–284.

Freytag, G., & MacEwan, E. J. (1895). Technique of the Drama: An Exposition of Dramatic Composition and Art; An Authorized Translation from the 6th German Edition. Griggs & Company, 1895.

García-Prieto, J., Bajo, R., & Pereda, E. (2017). Efficient computation of functional brain networks: toward real-time functional connectivity. Frontiers in Neuroinformatics, 11, 8.

Green, M. C., & Brock, T. C. (2000). The role of transportation in the persuasiveness of public narratives. Journal of Personality and Social Psychology, 79(5), 701–721. https://doi.org/10.1037/0022-3514.79.5.701

Haynes, J.-D. (2015). A primer on pattern-based approaches to fMRI: principles, pitfalls, and perspectives. Neuron, 87(2), 257–270.

Herman, D. (2009). Cognitive Narratology. In P. Hühn, J. Pier, W. Schmid, & J. Schönert (Eds.), Handbook of narratology (pp. 30–43). De Gruyter. https://doi.org/doi:10.1515/9783110217445

Hiskes, B., Hicks, M., Evola, S., Kincaid, C., & Breithaupt, F. (2022). Multiversionality: Considering multiple possibilities in the processing of narratives. Review of Philosophy and Psychology, 0123456789. https://doi.org/10.1007/s13164-022-00621-5

Huff, M., Meitz, T. G. K., & Papenmeier, F. (2014). Changes in situation models modulate processes of event perception in audiovisual narratives. Journal of Experimental Psychology: Learning, Memory, and Cognition, 40(5), 1377–1388. https://doi.org/10.1037/a0036780

Imhof, M. A., Schmälzle, R., Renner, B., & Schupp, H. T. (2020). Strong health messages increase audience brain coupling. NeuroImage, 216(August 2019), 116527. https://doi.org/10.1016/j.neuroimage.2020.116527

Ismail, L. E., & Karwowski, W. (2020). A graph theory-based modeling of functional brain connectivity based on eeg: A systematic review in the context of neuroergonomics. IEEE Access, 8, 155103–155135.

Khadem, A., & Hossein-Zadeh, G.-A. (2013). Comparing the robustness of brain connectivity measures to Volume Conduction artifact. 2013 20th Iranian Conference on Biomedical Engineering (ICBME), 209–214. https://doi.org/10.1109/ICBME.2013.6782220

Landauer, T. K., Foltz, P. W., & Laham, D. (1998). An introduction to latent semantic analysis. Discourse Processes, 25(2–3), 259–284.

Landauer, T. K., McNamara, D. S., Dennis, S., & Kintsch, W. (2013). Handbook of latent semantic analysis. Psychology Press.

Laurel, B. (1991). Computers as theatre reading. Mas: Addison-Wesley Publishing Company.

Lin, S.-F., Dale, K. R., McDonald, D. G., Collier, J. G., & Jones, K. (2019). Narrative Engagement and Vicarious Interaction with Multiple Characters. Mass Communication and Society, 22(3), 324–343. https://doi.org/10.1080/15205436.2018.1545034

Mitzdorf, U. (1985). Current source-density method and application in cat cerebral cortex: investigation of evoked potentials and EEG phenomena. Physiological Reviews, 65(1), 37–100. https://doi.org/10.1152/physrev.1985.65.1.37

Nastase, S. A., Gazzola, V., Hasson, U., & Keysers, C. (2019). Measuring shared responses across subjects using intersubject correlation. Social Cognitive and Affective Neuroscience, 14(6), 669–687. https://doi.org/10.1093/scan/nsz037

Nguyen, M., Vanderwal, T., & Hasson, U. (2019). Shared understanding of narratives is correlated with shared neural responses. NeuroImage, 184(September 2018), 161–170. https://doi.org/10.1016/j.neuroimage.2018.09.010

Petroni, A., Cohen, S. S., Ai, L., Langer, N., Henin, S., Vanderwal, T., Milham, M. P., & Parra, L. C. (2018). The Variability of Neural Responses to Naturalistic Videos Change with Age and Sex. Eneuro, 5(1), ENEURO.0244-17.2017. https://doi.org/10.1523/ENEURO.0244-17.2017

Poulsen, A. T., Kamronn, S., Dmochowski, J., Parra, L. C., & Hansen, L. K. (2017). EEG in the classroom: Synchronised neural recordings during video presentation. Scientific Reports, 7(1), 43916. https://doi.org/10.1038/srep43916

Rosenberg, M. D., Scheinost, D., Greene, A. S., Avery, E. W., Kwon, Y. H., Finn, E. S., Ramani, R., Qiu, M., Constable, R. T., & Chun, M. M. (2020). Functional connectivity predicts changes in attention observed across minutes, days, and months. Proceedings of the National Academy of Sciences, 117(7), 3797–3807.

Ryan, M.-L. (2007). Toward a definition of narrative. In D. Herman (Ed.), The Cambridge Companion to Narrative (pp. 22–36). Cambridge University Press. https://www.cambridge.org/core/product/identifier/CBO9781139001533A006/type/book_part

Sendi, M. S. E., Zendehrouh, E., Sui, J., Fu, Z., Zhi, D., Lv, L., Ma, X., Ke, Q., Li, X., & Wang, C. (2021). Abnormal dynamic functional network connectivity estimated from default mode network predicts symptom severity in major depressive disorder. Brain Connectivity, 11(10), 838–849.

Sendi, M. S. E., Zendehrouh, E., Turner, J. A., & Calhoun, V. D. (2021). Dynamic patterns within the default mode network in schizophrenia subgroups. 2021 43rd Annual International Conference of the IEEE Engineering in Medicine & Biology Society (EMBC), 1640–1643.

Shen, X., Finn, E. S., Scheinost, D., Rosenberg, M. D., Chun, M. M., Papademetris, X., & Constable, R. T. (2017). Using connectome-based predictive modeling to predict individual behavior from brain connectivity. Nature Protocols, 12(3), 506–518.

Sijmen Duineveld. (2022). No Title.

Silva, M., Baldassano, C., & Fuentemilla, L. (2019). Rapid memory reactivation at movie event boundaries promotes episodic encoding. BioRxiv, 39(43), 8538–8548. https://doi.org/10.1101/511782

Simony, E., Honey, C. J., Chen, J., Lositsky, O., Yeshurun, Y., Wiesel, A., & Hasson, U. (2016). Dynamic reconfiguration of the default mode network during narrative comprehension. Nature Communications, 7(1), 12141. https://doi.org/10.1038/ncomms12141

Song, H., Finn, E. S., & Rosenberg, M. D. (2021a). Neural signatures of attentional engagement during narratives and its consequences for event memory. Proceedings of the National Academy of Sciences, 118(33).

Song, H., Finn, E. S., & Rosenberg, M. D. (2021b). Neural signatures of attentional engagement during narratives and its consequences for event memory. Proceedings of the National Academy of Sciences, 118(33), e2021905118. https://doi.org/10.1073/pnas.2021905118

Song, H., Park, B., Park, H., & Shim, W. M. (2020). Cognitive and neural state dynamics of story comprehension. BioRxiv.

Song, H., & Rosenberg, M. D. (2021). Predicting attention across time and contexts with functional brain connectivity. Current Opinion in Behavioral Sciences, 40, 33–44.

Sonkusare, S., Breakspear, M., & Guo, C. (2019). Naturalistic Stimuli in Neuroscience: Critically Acclaimed. Trends in Cognitive Sciences, 23(8), 699–714. https://doi.org/10.1016/j.tics.2019.05.004

Vaccaro, A. G., Scott, B., Gimbel, S. I., & Kaplan, J. T. (2021). Functional Brain Connectivity During Narrative Processing Relates to Transportation and Story Influence. Frontiers in Human Neuroscience, 15(July), 1–14. https://doi.org/10.3389/fnhum.2021.665319

Varotsis, G. (2018). The plot-algorithm for problem-solving in narrative and dramatic writing. New Writing, 15(3), 333–347. https://doi.org/10.1080/14790726.2017.1374414

Zwaan, R. A. (1999). Situation Models. Current Directions in Psychological Science, 8(1), 15–18. https://doi.org/10.1111/1467-8721.00004

